# A CRISPR-associated Rossman-fold Ring Nuclease and Adenosine Deaminase Fusion Allosterically Converts ATP to ITP

**DOI:** 10.1101/2024.12.12.628194

**Authors:** Charlisa Whyms, Yu Zhao, Doreen Addo-Yobo, Huan He, Arthur Carl Whittington, Despoina Trasanidou, Carl Raymund P. Salazar, Raymond H.J. Staals, Hong Li

## Abstract

The recently identified CARF (CRISPR-associated Rossman-fold) family of proteins play a critical role in prokaryotic defense, mediating cOA (cyclic oligoadenylate)-stimulated ancillary immune responses in the type III CRISPR-Cas systems. Whereas most previously characterized CARF proteins contain nucleic acids or protein degradation effectors, a subset comprises <u>C</u>ARF-fused <u>a</u>denosine <u>d</u>eaminase (ADA) (Cad1) and has a yet to be determined function. Here we present biochemical and structural analyses of a ring nuclease Cad1, revealing its unexpected role in deaminating adenosine-5′-triphosphate to inosine-5′-triphosphate in a cOA-dependent manner. Despite an overall structural similarity to canonical ADA enzymes, the ADA domain of Cad1 possesses unique structural features underlying its specificity for ATP. Supported by mutational analysis, we demonstrate an allosteric link between the cOA-binding CARF and the ADA domain, suggesting that Cad1 is a cOA-stimulated effector that influences cellular metabolic processes.

**Highlights:**

- TaqCad1 converts ATP to ITP in a magnesium- and cOA-dependent manner
- Cryo-EM structures reveal TaqCad1 forms a homohexamer
- Cryo-EM structures reveal how cA_4_ is degraded and modulates the ADA active site

## Introduction

Virtually all domains of life utilize linear or cyclic nucleotide-based second messengers to regulate diverse cellular processes in response to external or internal stimuli ^1^. Recently, the cyclic oligonucleotide (cO)-based signaling strategy has been expanding to prokaryotes in the CRISPR (Clustered Regularly Interspaced Short Palindromic Repeats)-Cas (CRISPR-associated) systems. CRISPR-Cas confers the adaptive immunity to their hosts against the viruses and other invasive mobile genetic elements ^2-4^. The Type III CRISPR-Cas systems exemplify this functionality by eliciting multi-pronged and multi-leveled anti-viral responses. The Type III CRISPR-Cas effector complex senses, via a protein-associated CRISPR RNA (guide RNA), and degrades the viral transcripts (target RNA), which triggers two enzymatic activities within its Cas10 subunit: the nonspecific shredding of single stranded DNA and the synthesis of cOA molecules from adenosine triphosphate (ATP) ^5,6^. The cOA molecules subsequently stimulate the unique CARF (CRISPR-associated Rossman-fold) domain-containing family of proteins that further assist with clearing the infection. An analogous process occurs in the CBASS (cyclic-oligonucleotide-based-anti-phage-signaling-system) systems in which phage infection also stimulates the production of diverse cOAs that then activate downstream proteins, ultimately halting the propagation of the phages ^7-9^. The cOA-mediated signaling processes provide a wealth of insights into enzyme regulation principles that benefit both basic research and biotechnology development.

Bioinformatic analysis has unveiled an impressive collection of diverse CARF proteins that are made of the CARF domain alone or a fused effector domain of various enzymatic function ^10^. The CARF family of proteins have been classified into 9 major and 13 minor clades ^11^. The most common CARF proteins associated with the Type III CRISPR-Cas systems are the CARF-fused HEPN (Higher Eukaryotes and Prokaryotes nucleotide-binding) or PD/D/ExK nucleases such as Csm6/Csx1 and the CARD1 like proteins ^5,6,12-14^. Structural and biochemical analysis reveal that these CARF-nucleases bind specific cOA molecules that allosterically enhance the catalytic activity of their fused effector domains ^9^ by diverse mechanisms ^15,16^, ranging from controlling the catalytic metal ions ^12,17^ to oligomerization ^18^ of the effector domains. While the activation of CARF-nucleases by cOA is pivotal for Type III immunity, this process can potentially lead to cell toxicity, thereby necessitating the requirement for regulatory mechanisms. Consequently, certain organisms possess CARF proteins that, with specific amino acid substitutions near the cOA binding site, function as either independent ring or self-limiting nucleases to remove cOA molecules ^19-24^.

In contrast to the well-characterized CARF nucleases, little is known about <u>C</u>A<u>R</u>F-fused adenosine <u>d</u>eaminases (Cad1), members of the CARF5 clade group. These CARF proteins possess the characteristics indicative of cOA binding and/or cleavage activities ^11^, suggesting the possible cOA-regulated deamination functionality. Adenosine deaminases (ADA) are enzymes that convert adenosine and deoxyadenosine (A) to inosine and deoxyinosine (I), respectively ^25^. Infection-triggered A-to-I conversion could potentially deplete the pool of (deoxy)adenosine required for viral replication and thus virulency. Here we report biochemical and structural characterization of the predicted Cad1 from *Thermoanaerobaculum aquaticum* (TaqCad1) that is distinct from the CARF proteins characterized to date. The CRISPR locus of the *T. aquaticum* genome encodes the known components for spacer integration, the Cas proteins constituting the Type III-B CRISR-Cas effector complex, ancillary nuclease Csx1, and Cad1 (Figure 1a). We show that Cad1 binds specifically to and acts as a ring nuclease for the cyclic tetra-adenosine (cA_4_) in regulation of the nuclease activity of Csx1. Surprisingly, the ADA domain of Cad1 does not convert adenosine to inosine, but instead, the adenosine triphosphate (ATP) to inosine triphosphate (ITP). Supported by biochemical data, the cryoEM structures of TaqCad1 in the presence of cA_4_ reveal an hexameric architecture and how the two domains interact with the ligand. Our findings suggest a novel function of this subset of the CARF family in defending viruses, potentially by impacting metabolic functionality in cells.

**Figure 1.**
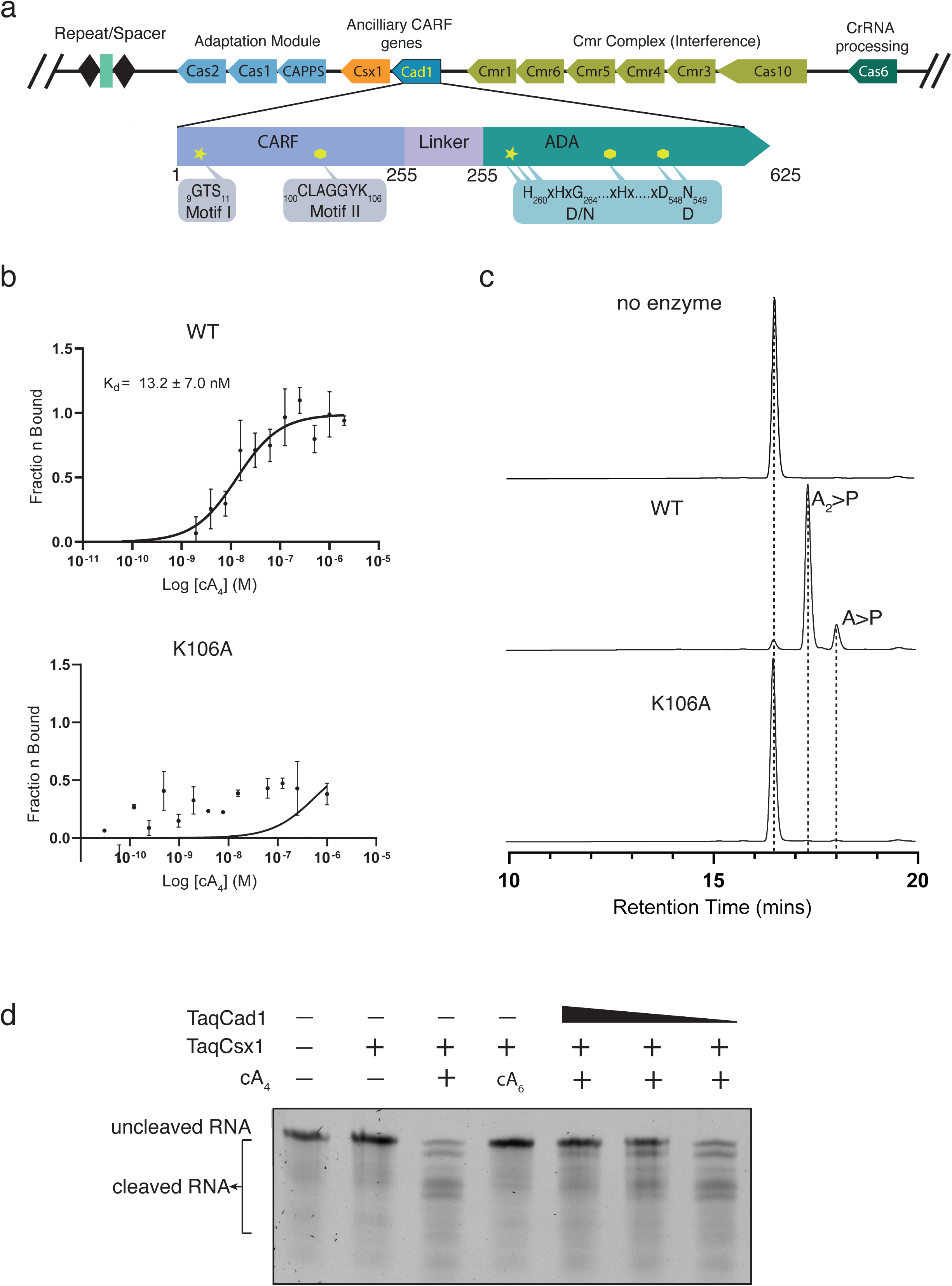
TaqCad1 features and biochemical activities. **a.** *Thermoanaerobaculum aquaticum* CRISPR locus and TaqCad1 sequence features. Key amino acids along with their residue numbers for the two enzymatic domains are shown in the insets where “D/N” and “D” describe the residues conserved in canonical adenosine deaminase active sites. **b.** Microscale thermophoresis (MST) titration curves of cyclic tetra-adenosine (cA_4_) to TaqCad1 or its mutant K106A. The apparent *K_d_* is estimated by fitting the binding isotherm to a one-specific binding model. Three experimental replicas were formed from which standard deviation was obtained. No data fitting was attempted for K106A titration. **c.** Elution profiles of cA_4_ and its hydrolyzed products on HPLC by TaqCad1 or K106A. **d.** The impact of TaqCad1 ring nuclease activity on TaqCsx1 RNA cleavage activity.

## Results

### TaqCad1 Binds and Hydrolyzes cA_4_

The CARF domain of TaqCad1 contains characteristic motifs for binding cOA molecules (Figure 1a). Motif-II residues are crucial for cOA binding, while motif-I residues are largely responsible for cOA degradation. To confirm this activity and to distinguish the types of cOA molecules it interacts with, we employed microscale thermophoresis (MST) in measuring binding dissociation constants of the recombinantly expressed TaqCad1 and its variants (Figure 1b & Supplementary Figure 1a). Titration of 0.1 nM – 4 μM cA_4_ to 50 nM fluorophore-labeled TaqCad1 revealed an apparent disassociation constant of 13.2 +/- 7.0 nM (Figure 1b), similar to those measured for cA_4_ binding to Sso2081 (Crn1) ^21^ or Sis0811 ^26^ CARF proteins. In contrast, MST titration of cA_6_ with TaqCad1 resulted in an estimated apparent dissociation constant of ∼1.4 µM (Supplementary Figure 1b), suggesting that cA_4_ is the preferred ligand. Furthermore, mutation of the motif II residue, Lys106, to alanine (K106A) largely abolished binding of cA_4_ (Figure 1b).

To evaluate if TaqCad1 possesses the ring nuclease activity, we incubated it with cA_4_ or cA_6_ for one hour at 37°C and detected the reaction products by High Performance Liquid Chromatography (HPLC) and mass spectroscopy. Whereas cA_4_ was readily degraded (Figure 1c), cA_6_ remained intact (Supplementary Figure 1c). The products of degradation are predominately di-adenosine cyclic phosphate (A_2_>P) and some adenosine cyclic phosphate (A>p) (Supplementary Figure 1d). Furthermore, K106A did not degrade cA_4_, consistent with its impaired binding to cA_4_ (Figure 1c). As shown for other ring nucleases, cA_4_ degradation by TaqCad1 did not require any metal, suggesting a general acid-base mechanism resembling that of RNase A ^27^. Finally, the ring nuclease activity of TaqCad1 functions as a likely regulator of the co-expressed TaqCsx1 by inhibiting its cA_4_-dependent ribonuclease activity (Figure 1d).

### TaqCad1 Lacks Adenosine Deamination Activity

The high sequence homology of the ADA domain of TaqCad1 to the known adenosine deaminases prompted us to investigate whether it promotes the conversion of A-to-I. For this, we made use of the characteristic change in UV absorption from adenosine at 265 nm to inosine at 235 nm. However, we were not able to detect any A-to-I activities, either in the presence or absence of cA_4_ (Figure 2a). We also employed mass spectrometry to detect the possible A-to-I conversion under the similar condition used for the spectroscopic method. We found that the wild-type TaqCad1 exhibits a weak A-to-I conversion (Figure 2a).

**Figure 2.**
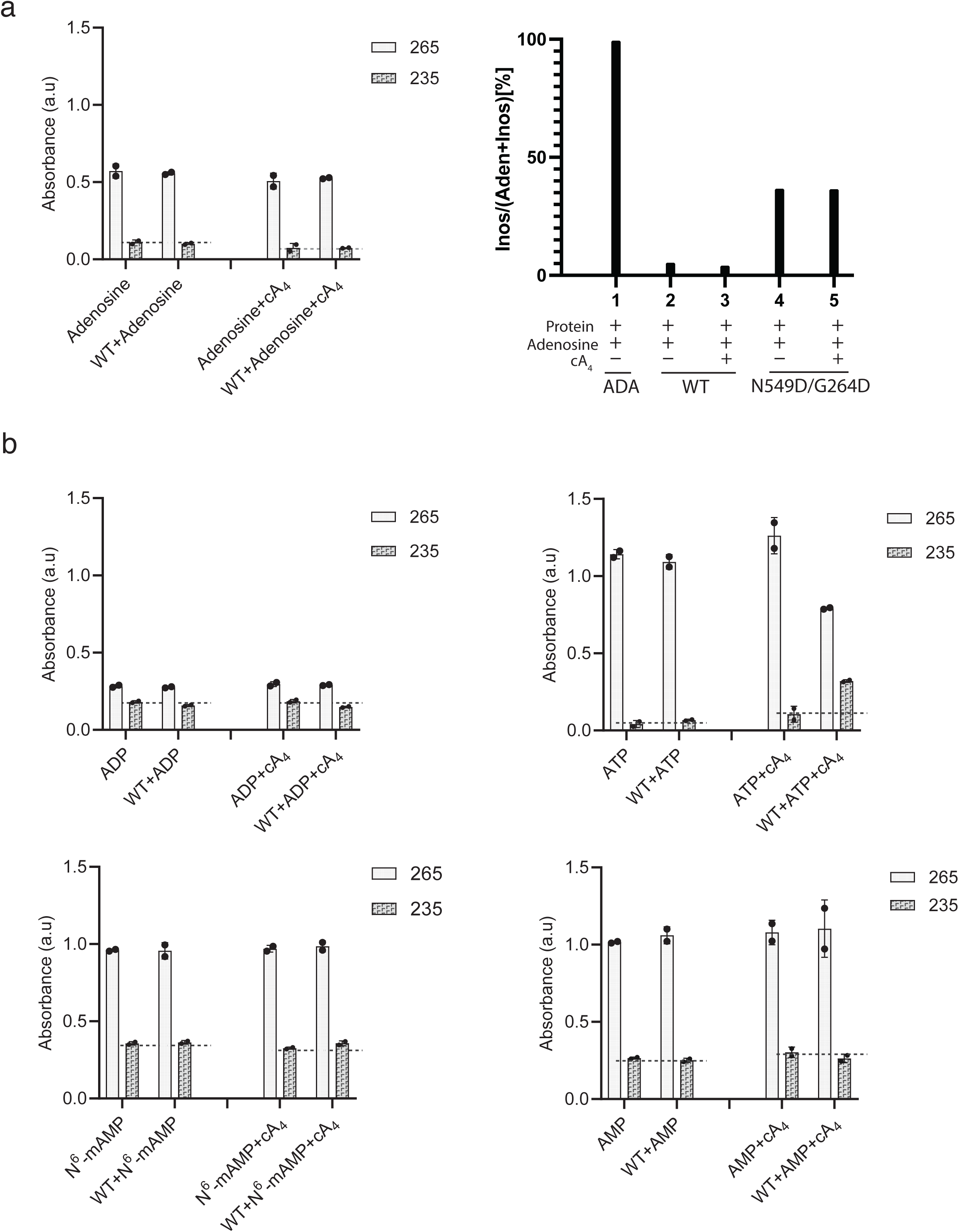
UV absorbance and mass spectrometry analysis of TaqCad1 deamination activities. **a.** Detection of spectroscopic or mass changes by using TaqCad1 wildtype (WT) with adenosine. Left, UV absorbance. Right, the products detected by mass spectroscopy. “ADA” denotes the product by a control adenosine deaminase. **b.** Detection of spectroscopic changes by using TaqCad1 wildtype (WT) with ADP, ATP, AMP, and N^6^-methyl-AMP (N^6^-mAMP). UV absorbance at wavelength 265 and 235 measures the characteristic absorbance of adenosine and inosine, respectively. TaqCad1-mediated reactions were dephosphorylated by calf intestinal alkaline phosphatase following heat deactivation.

Notably, the putative active sites of TaqCad1 and its close homologs contain two substitutions from the canonical ADA enzymes, a catalytic aspartate (Asp296 in mouse ADA) ^28^ by asparagine (Asn549 in TaqCad1) and a substrate-binding aspartate (Asp19 in mouse ADA) ^28^ by glycine (Gly264 in TaqCad1) (Figure 1a). Previous studies showed that mutation of the Asp549-equivalent aspartate, Asp296, to asparagine in mouse ADA severely impaired substrate binding and catalytic efficiency ^28^. Furthermore, the carboxylate of Asp19 of mouse ADA forms two hydrogen bonds with the bound adenosine, one to its 5’-hydroxyl and another to its 3’-hydroxyl oxygen, respectively ^29^. Substitution of both residues in TaqCad1 could thus render it catalytically inactive. However, replacing these residues with the canonical ADA residues (N549D/G264D) failed to rescue the low deamination activity (Figure 2a), indicating that these residues are not the sole or primary factors contributing to the lack of adenosine deamination activity.

### Cad1 Binds and Deaminates ATP

We expanded our search for substrate to other metabolites that contain adenosine moiety by measuring their UV absorbance changes. We included AMP, ADP, ATP, N^6^-methyl-AMP that are known substrates for related adenosine deaminases (Figure 2b). Among these, ATP exhibited significantly strong conversion of inosine absorbance only in the presence of cA_4_ (Figure 2b), suggesting the possibility that it may be the substrate for TaqCad1, especially given the previous example of ATP to ITP conversion by the deaminase of the anti-phage system, RADAR ^30,31^.

We further confirmed deamination of ATP by TaqCad1 by analyzing the reaction products by High Performance Liquid Chromatography (HPLC). Indeed, in presence of either cA_4_ or cA_6_, as well as a divalent ion (Mg^2+^ or Ca^2+^) ATP was readily converted to ITP (Figure 3a) while AMP, ADP or N^6^-methyl-AMP were not (Supplementary Figure 2a-2c). The deamination activity requires Mg^2+^, Ca^2+^ or Zn^2+^, with both Ca^2+^ and Zn^2+^ showing weaker efficacy (Figure 3b). Interestingly, the N549D or the N549D/G264D variant of TaqCad1 did not react to ATP (Figure 3c & Supplementary Figure 2d), suggesting a distinct active site of TaqCad1 from the canonical adenosine deaminases. Furthermore, the ring-nuclease variant, K106A, also failed to convert ATP (Figure 3b & Supplementary Figure 2d), consistent with its lack of binding to cA_4_ (Figure 1c). Our data suggest that TaqCad1 is distinct from canonical adenosine deaminases and specific for ATP.

**Figure 3.**
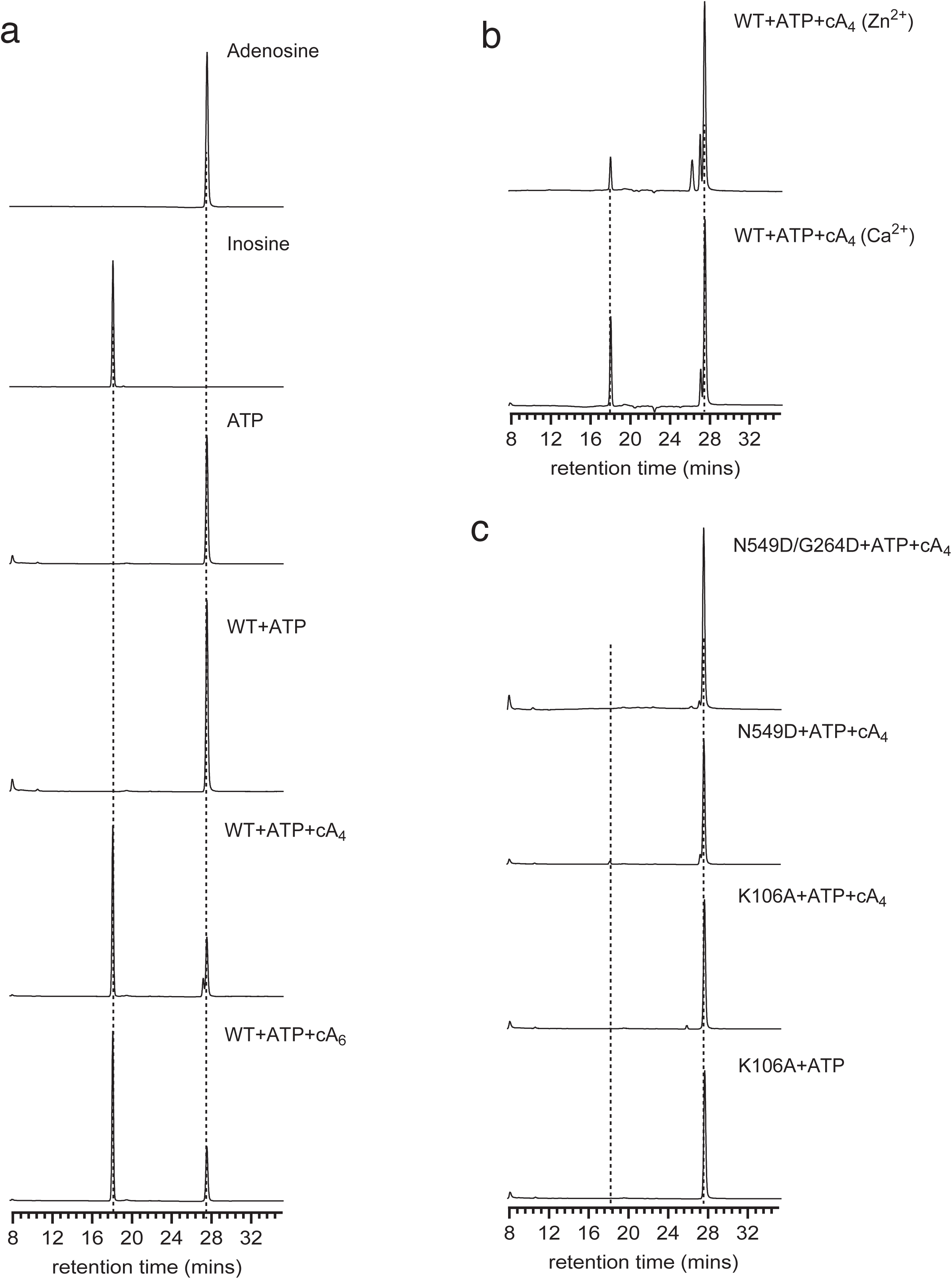
High Performance Liquid Chromatography (HPLC) analysis of the possible products of ATP treated with TaqCad1 or its variants. a. HPLC elution profiles of ATP reacted with TaqCad1 in the presence or absence of cA_4_ or cA_6_. b. HPLC elution profiles of ATP reacted with TaqCad1 in the presence of cA4 and Ca^2+^ or Zn^2+^. c. HPLC elution profiles of ATP reacted with TaqCad1 variants in the presence of cA4 and Ca^2+^ or Zn^2+^.

### TaqCad1 Forms a Homohexamer that Binds Three cA_4_

To learn the structural basis, we obtained cryoEM structures of the wild-type TaqCad1 without a ligand or in the presence of cA_4_, resulting in an apo and a cA_4_-bound structures (Supplementary Figures 3, Supplementary 4 & Supplementary Table 1). In both cases, TaqCad1 forms a hexameric structure with a pseudo-D3 symmetry, composed of three homodimers (Figure 4). The ADA domains of the six subunits create a donut-shaped core, while the CARF domains extend outward in a propeller-like configuration (Figure 4). In both complexes, a coordinated metal ion is observed at the conserved active site of the ADA domain (Figure 4b), despite that no divalent ions were added during structural determination. Given the similarity to the canonical zinc-containing ADA enzymes, we modeled the ion as Zn^2+^. In the cA_4_-bound complex, two cleaved A_2_>P molecules are observed in the CARF domain (Figure 4b & Supplementary Figure 5).

**Figure 4.**
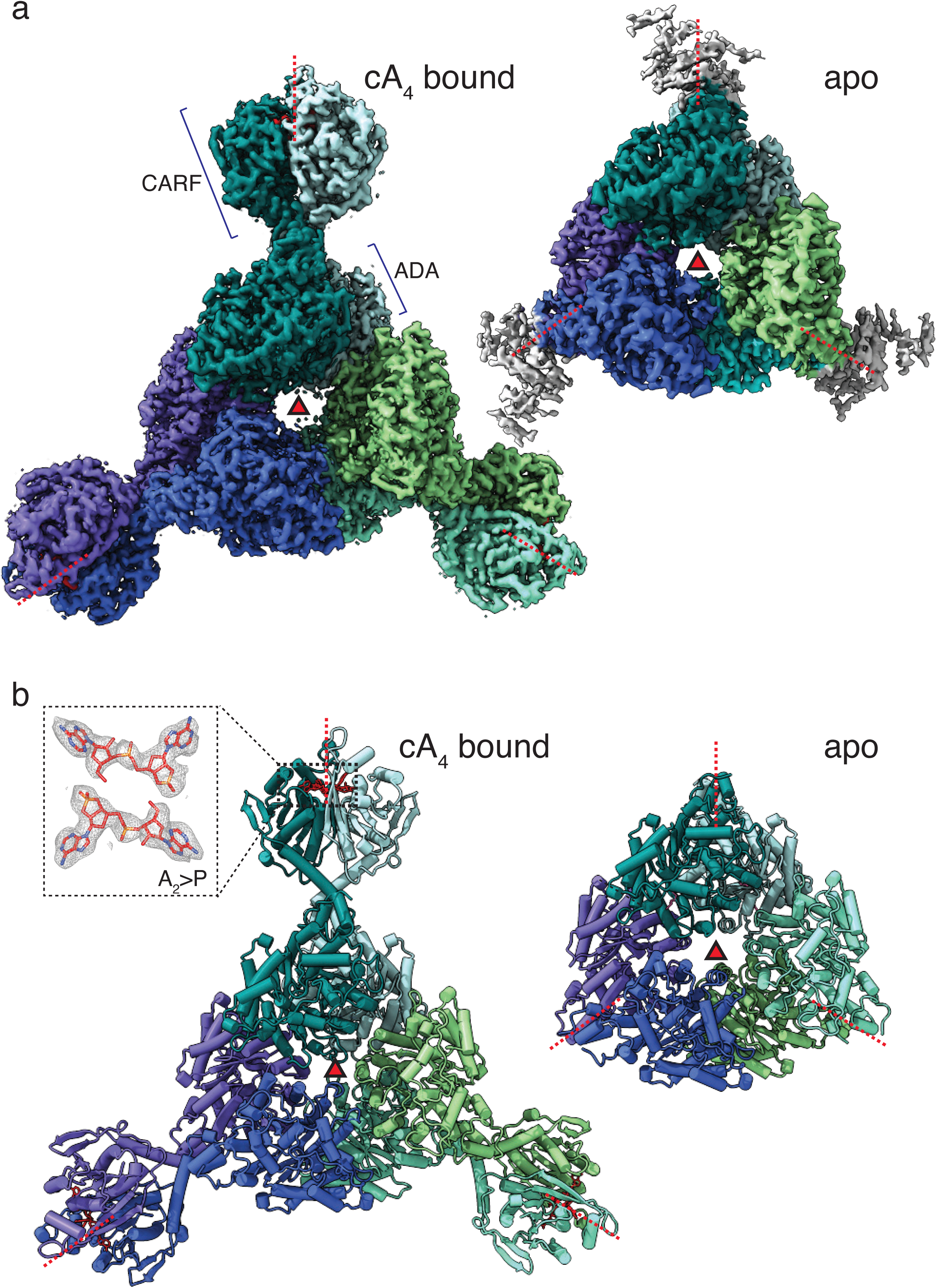
Structure overview and detailed ligand binding of TaqCad1. The pseudo-3-fold axis is indicated by filled solid triangle while the three perpendicular 2-fold axes are indicated by dashed lines in red. **a.** Structure of the TaqCad1 hexamer bound with cA_4_. Each protomer is assigned a unique color. Top, cryoEM density. Bottom, cartoon model. Inset, close-up view of the density around one bound A_2_>P, the primary hydrolyzation product of cA_4_. The pseudo-3-fold axis is indicated by filled triangle while the three perpendicular 2-fold axes are indicated by dashed red lines. **b.** Structure of the Apo TaqCad1 Top, cryoEM density. Bottom, cartoon model. Note that the density for the CARF domain is not modeled due to the heterogenous and weak density.

There is significant conformational heterogeneity in the CARF domain, especially when cA_4_ is not present. Even when cA_4_ is bound, the CARF domains within the same hexameric particles exhibit alternative conformations. We, therefore, reconstructed the cA_4_-bound dimer structure using C3 symmetry-expanded particles (good dimers) to yield the best possible maps, leading to 2.4 Å and 2.3 Å resolution for the CARF and ADA domain, respectively (Supplementary Figure 3 & Supplementary Table 1). The same procedure did not improve the density of the CARF domain in the complex without cA_4_. The density of the hexameric apo complex is obtained by applying C3 symmetry of the particles containing an intact hexamer (good hexamers) during reconstruction (Supplementary Figure 4 & Supplementary Table 1). The CARF domain in the apo structure is largely disordered (Figure 4), suggesting a role of cA_4_ in rigidifying the CARF domain.

The homodimer is similarly formed as other CARF family of proteins by using its Rossmann fold core (2^nd^ P-loop and α6, residues 105-120) and the helix-turn-helix insertion (residues 211-229) (Figure 5a & Supplementary Figure 5). The dimer interface buries an extensive solvent accessible area (∼1800 Å^2^), suggesting that it is the primary interface of the assembly. Notably, the two ADA domains of each dimer do not engage in close contacts. Rather, they independently dimerize with the ADA domain of two different neighboring dimers with ∼900 Å^2^ buried surface area, providing the structural basis for the assembly of the three pairs of homodimers (Figure 5a). The helix ⍺21 within the ADA domain is the major secondary element involved in the ADA-ADA contacts that comprise of a mix of hydrophobic and polar interactions. Consistently, mutation of Arg408 and Ala4ll to Asp and Glu (R408D/A411E), respectively, disrupted the hexameric assembly as revealed by the altered size-exclusion elution profile (Figure 5b). Interestingly, the disassembled dimer, while remains active in cleaving cA4 (Figure 5c), can no longer convert ATP to ITP (Figure 5d). This result suggests that hexamerization is only required for deamination.

**Figure 5.**
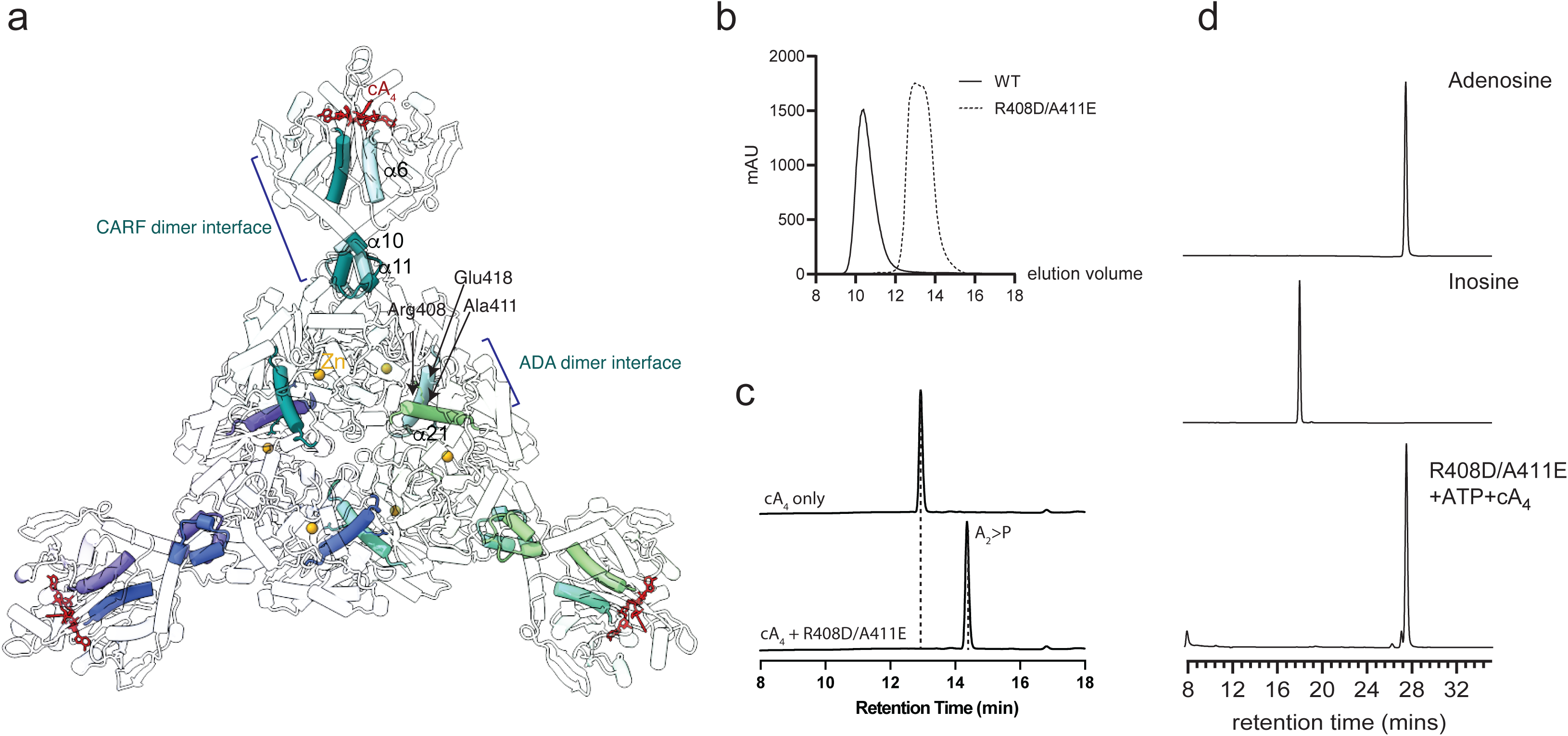
Key features of TaqCad1 oligomerization. **a.** Highlight of the dimerization and hexamerization interfaces mediated by the CARF and ADA domain, respectively. Helices α6, α10, and α11 mediate dimerization while α21 mediates hexamerization. These helices are shown as solid cylinders and labeled. The bound cA_4_ molecules are shown as red stick models. Key residues of α21 involved in hexamerization are labeled. **b.** Gel filtration elution profiles of the wild-type (WT) and R408D/A411E variant (R408D/A411E) indicating significantly reduced hydrodynamic radius upon mutation. **c.** HPLC profiles of the digested cA_4_ molecule by the R408D/A411E mutant (bottom) in comparison with cA_4_ alone (top). **d.** HPLC analysis of ATP deamination by the wild-type (WT) and the R408D/A411E mutant in the presence of cA_4_.

The cA_4_-bound structure shows that one cA_4_ molecule is bound at the interface of two CARF domains, leading to three cA_4_ bound per hexamer (Figure 4 & Supplementary Figure 5a). The discontinued density suggests that the bound cA_4_ is linearized with a 2′, 3′ cyclic phosphate and a 5′-OH termini at the two cleavage sites, creating two A_2_>P products (Figure 4 & Supplementary Figure 5a). Both Lys106 residues of the dimer are placed near the scissile phosphate with the amino nitrogen close to the leaving 5’-hydroxyl oxygen (∼3.2 Å) (Supplementary Figure 5a), consistent with the detrimental effect upon its mutation (Figure 1b). In addition, Thr10 and Ser11 have close contacts with reactive groups and suggest a general base mechanism in which Thr10 likely deprotonates the nucleophile and Lys106 protonates the leaving 5’-hydroxyl or, with Ser11, stabilizes the negatively charged pentavalent intermediate (Supplementary Figure 5a). The fact that cA_4_ is cleaved by composite catalytic centers made of residues from both CARF domains suggests an importance of dimerization to cA_4_ binding and degradation.

### cA_4_ Modulates the ADA Active Site

To learn the structural basis for cOA-mediated activation of the deaminase activity, we compared the cA_4_-bound structure to the apo structure. In the absence of cA_4_, the CARF domain is significantly more flexible (Figure 4 & Figure 6). In addition, the opening of the donut-shaped ring is enlarged slightly by ∼ 5 Å, primarily through a loop-to-helix transition of the region 435-454 (helix α22 in the cA_4_-bound structure) (Figure 6a & Supplementary Figure 5b). The loop portion of this region in the apo structure restructured to the more rigid α22, leading to a slight closure of the central ring when cA_4_ is bound (Figure 6a). Notably, the restructured α22 is adjacent to the catalytic center, especially the metal coordination ligand His461. When the ADA domain of the cA_4_-bound structure is superimposed on that of the apo structure, an upward shift in the bound Zn^2+^ and its coordination ligands His461 and Asp548 is observed (Figure 6b), which likely aligns the catalytic groups for catalysis.

**Figure 6.**
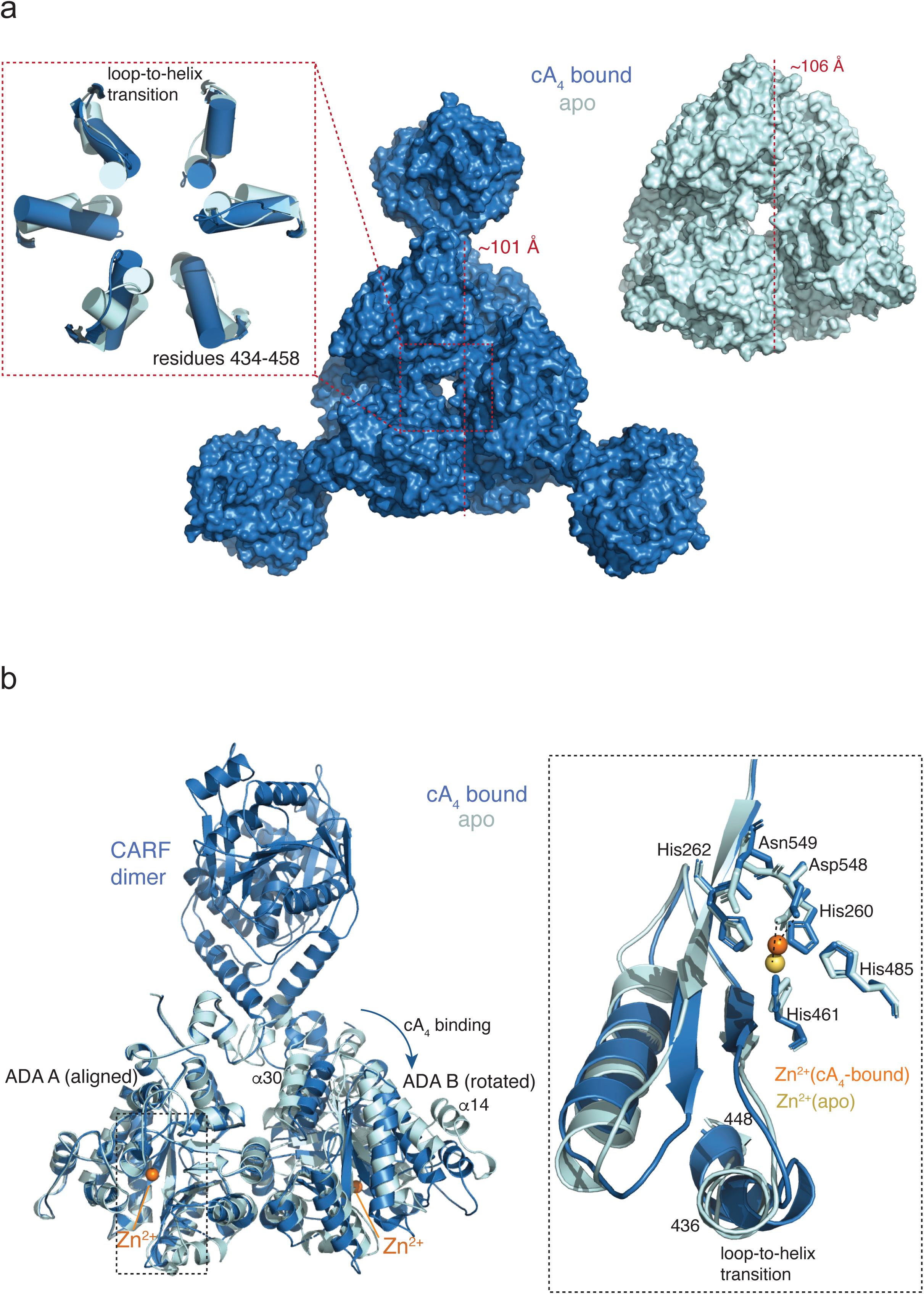
Molecular mechanism of TaqCad1 activation by cA_4_. a. Comparison of overall structures of TaqCad1 in the presence (blue) and absence (light cyan) of cA_4_. Inset displays close-up view of the center of the ADA ring. b. Comparison of the ADA active site in the presence (blue) and absence (light cyan) of cA_4_. One of the ADA domains is aligned between the cA_4_-bound and the cA_4_-free CARF dimer (ADA A). The observed rotation of the other ADA domain (ADA B) is indicated. Inset displays close-up view of the ADA active site in comparison with key residues labeled.

Though the ADA domain of TaqCad1 has an overall similarity with canonical adenosine deaminases (Supplementary Figure 6), the immediate surroundings of the substrate binding pocket differ substantially (Figure 5a). We thus assessed the importance of several residues that may potentially be involved in binding the substrate ATP. Indeed, mutation of Met311 to aspartate (M311D), Gly314 to phenylalanine (G314F), Asn549 to aspartate (N549D) are all detrimental to the ATP to ITP conversion activity (Supplementary Figure 2), suggesting the active site of TaqCad1is evolved to be specific for ATP.

### TaqCad1 Has a Mild Defense Activity at 37°C

Most CARF domain-containing proteins are involved in some defense function, and their activation often leads to either cell dormancy or abortive infection ^32^. Here, we investigated whether the enzymatic activity of TaqCad1 on ATP causes decreased cell viability. Therefore, we co-expressed TaqCad1 and a cA_4_-producing type III complex from *Treponema succinifaciens* in *E. coli* BL21-AI. As controls, we also co-expressed a cA_4_-responsive *T. succinifaciens* Card1 (TsuCard1) and an empty vector with the *T. succinifaciens* type III complex as positive and negative controls, respectively ^12^ (Supplementary Figure 7).

Target and non-target plasmids were used to express IPTG-inducible protospacers that were either complementary or non-complementary to the spacer, respectively ^33^. These were electroporated into the abovementioned strains, and ten-fold serial dilutions were plated onto IPTG-containing plates to induce the production of target or non-target RNA. When TaqCad1 is present, the transformation efficiency of the target plasmid was significantly lower by a modest one order of magnitude compared to the non-target plasmid (Figure 7a). Interestingly, the presence of TaqCad1 and target RNA led to visibly smaller colonies as compared to the non-target condition, suggesting slower growth rates (Figure 7b).

**Figure 7.**
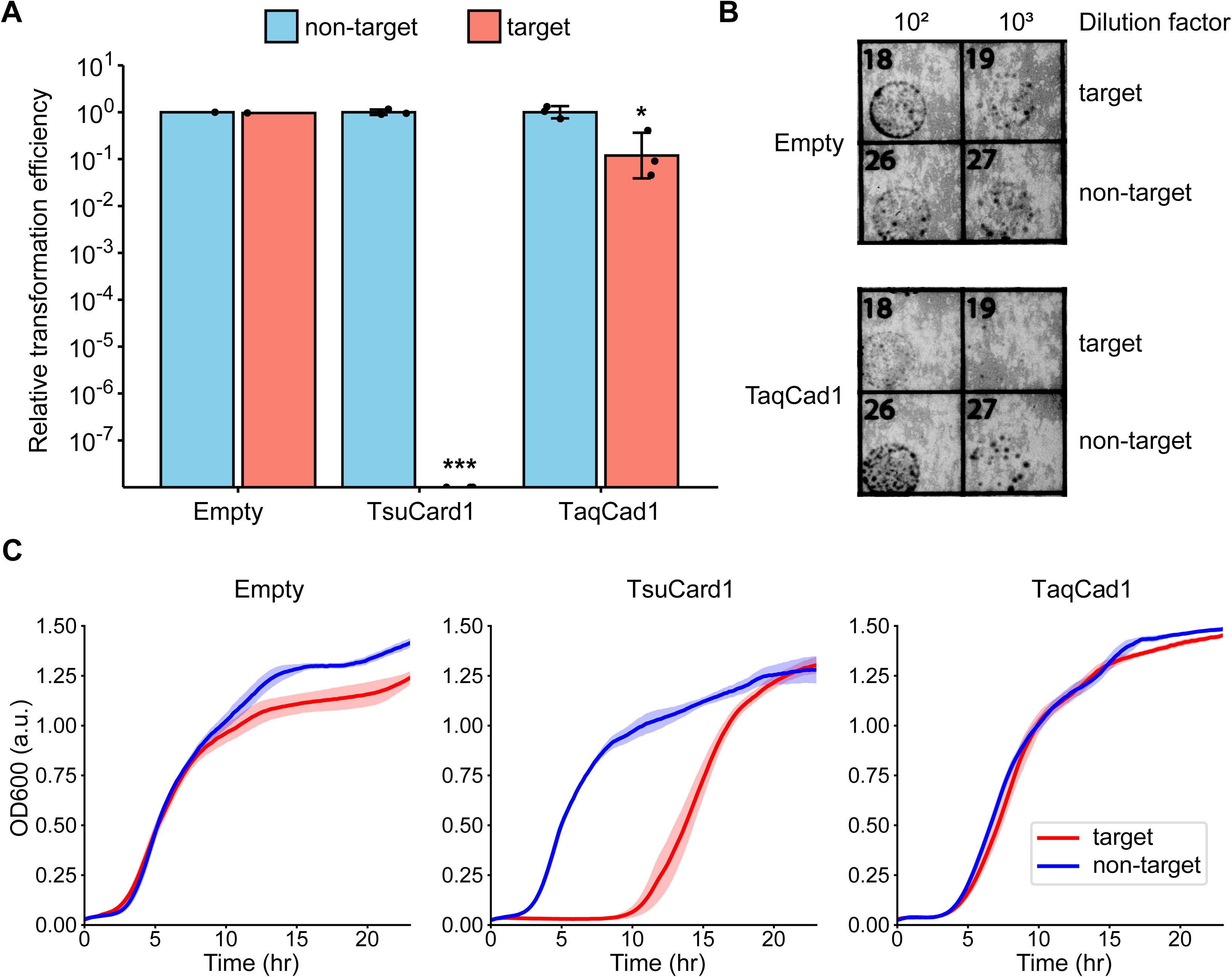
*In vivo* plasmid challenge assays to investigate the phenotype TaqCad1 activation. **a.** Transformation efficiencies (relative to non-target control) of target and non-target plasmids in *E. coli* co-expressing TaqCad1 and cA_4_-producing *Treponema succinifaciens* type III CRISPR-Cas complex. Error bars represent the standard deviation of the mean. Statistical significance was calculated using Welch’s t-test (n = 3 biological replicates). * *p* < 0.05, *** *p* < 0.005. **b.** Comparison of colony sizes obtained from the assay after overnight growth. **c.** Growth of transformants in liquid culture, measured as the optical density at 600 nm (OD_600_). Data represent the mean ± standard deviation of the mean (n = 3 biological replicates).

We also tracked the growth of transformants in liquid cultures, to better understand the effect of TaqCad1 activation on growth rate. While the presence of TaqCad1 does lead to a longer lag phase compared to the empty vector control, the presence or absence of target RNA did not have cause a difference in the lag phase (Figure 7c). These suggest that TaqCad1 has a minimal effect on growth. However, it should be noted that the assays were done at 37°C, well below the expected optimal temperature of the protein (60°C). This, in addition to the ring nuclease activity of TaqCad1, may have minimized the cA_4_-dependent defense activity of the protein. Altogether, these suggest that TaqCad1 activation via cA_4_ production upon target RNA detection has a mild defense response at 37°C.

## Discussions

We provide evidence that an adenosine deaminase-like enzyme fused with a ring nuclease CARF apparently has the ability to convert ATP to ITP in a cOA-dependent manner. In addition, the ring nuclease of TaqCad1 regulates the RNA shredding activity of another co-functional CARF nuclease, Csx1. Our structural analysis shows that the CARF domain forms the canonical cA_4_-binding and degradation module whereas the ADA-like domain both mediates oligomerization and interacts with ATP. Thus, TaqCad1 is a dual functional enzyme mediated by cOAs, both as a ring nuclease in regulating Csx1 and itself and as a deaminase to accumulate the level of ITP. Despite the robust enzymatic activities in vitro, interestingly, an in vivo plasmid challenge assay revealed a mild defense activity.

Intriguingly, a recent study identified that the anti-phage system, RADAR, possesses an adenosine deaminase subunit, RdrB that also converts ATP or dATP to ITP or dITP ^30,31^. It was suggested that the accumulation of ITP or dITP achieves anti-phage activity by both influencing the nucleotide pool and/or nucleic acid synthesis ^31^. TaqCad1 may exert the same cellular effects upon activation by cOAs, although its efficient ring nuclease activity may prevent a long-lasting effect from being detected at a significant level. Interestingly, whereas RADAR is able to deaminate a number of adenosine-containing metabolites, TaqCad1 is specific for ATP, which highlights their structural differences (Supplementary Figure 6).

While CARF-proteins primarily exist as homodimers ^12,13,17,22^, some instances show their presence as a monomer ^34^ and hexamer ^18^. Regardless of the oligomeric configuration, these proteins detect cOA activations through the linkage from the CARF to the effector domain. TaqCad1 forms a hexamer capable of binding three cA_4_ or cA_6_. The fact that ATP-to-ITP conversion is only observed upon cA_4_ or cA_6_ binding suggests an allosteric regulation of ATP binding by the second messenger produced upon viral infection. Consistently, we observed an overall reduction of the donut-shaped ring size formed by the ADA domains when cA_4_ is bound to the CARF domain. As a result, we observed a noticeable restructuring of the ADA active site. This is likely the structural basis for cA_4_-mediated regulation of ATP conversion of the ADA domain.

## Materials and Methods

### Cloning, Protein Expression and Purification

The complete TaqCad1 sequence (1–625) was cloned into a pET28a vector, using NcoI and BamHI restriction sites, giving rise to TaqCad1 fused with a C-terminal hexa-histidine tag. Mutations of the TaqCad1 were introduced via Q5 mutagenesis (New England Biolabs) and confirmed by full plasmid sequencing (Eurofins Genomics). For over expression, the protein-coding plasmids were transformed into NiCo21 (DE3) cells (New England Biolabs) that were then cultured at 37 °C to an OD_600_ of 0.8 and induced with 0.3 mM isopropyl β-D-1-thiogalactopyranoside (IPTG) at 18 °C overnight. Cells were harvested by centrifugation and stored at −80 °C. Defrosted cell pellets were lysed via sonication in a lysis buffer comprising 20 mM HEPES at pH 7.5, 200 mM NaCl, 50 mM imidazole, 5% glycerol, and 5 mM phenylmethylsulfonyl fluoride (PMSF). After centrifugation at 16,000 RPM for 30 minutes, the supernatant was applied onto Nickel-NTA Resin (GE Healthcare), followed by extensive washing with the lysis buffer with 50 mM imidazole. The target protein was eluted using the lysis buffer with 500 mM imidazole. The elutant was loaded onto the Mono Q® 5/50 GL (Cytiva) column and eluted with a linear gradient from 100 mM to 1 M NaCl over 45 minutes. The peak fractions were collected and concentrated before being loaded onto the Superdex 200 16/60 column pre-equilibrated in the gel filtration buffer (50 mM HEPES pH 7.5, 150 mM NaCl, 5 mM ßME). The target proteins were collected and flash-frozen in liquid nitrogen for storage at −80 °C.

The complete TaqCsx1 sequence (1–458) was amplified from the *Thermoanaerobaculum aquaticum* genomic DNA and cloned into pACYduet vector with a C-terminal hexa-histidine tag. TaqCsx1 was expressed in E*. coli* NICO cells and purified similarly as TaqCad1 described above.

### Cryo-EM sample preparation, data collection, and 3D reconstruction

The cA_4_-free or cA_4_-bound complexes were prepared with TaqCad1 at approximately 3 mg/mL (∼43 μM) concentration without or with cA_4_ at 105 μM in a buffer composed of 20 mM HEPES at pH 7.5, 150 mM NaCl, and 5 mM ß-mercaptoethanol (ßME) at room temperature for 15 minutes. Subsequently, a 4 μL volume of each solution was applied onto UltrAuFoil 300 mesh R1.2/1.3 grids (Quantifoil) glow-discharged with Gatan Solarus plasma cleaner 950. After 30 seconds wait in the Vitrobot (FEI Mark IV) chamber under 100% humidity and 18C, grids were blotted for 3 seconds and flash frozen in liquid ethane. The prepared grids were then stored in liquid nitrogen until data collection.

The raw micrographs of both data sets were collected at the Laboratory for BioMolecular Structure (LBMS) of the Brookhaven National Laboratory using a Krios G3i cryo transmission electron microscope equipped with a Gatan K3 direct electron detector (ThermoFisher Scientific). Movies were recorded at a nominal magnification of 81,000 in a super-resolution mode with an energy filter of 15 eV, corresponding to a corrected physical pixel size of 0.53 Å/pixel. A total dose of 60 e^-^/ Å^2^ was applied over 60 frames with random defocus set to -0.8 to -2.5 µm. Motion correction was performed in bin 2 mode using MotionCorr2 and the contrast transfer function (CTF) estimation was carried out using Gctf ^35^. A total of 8597 micrographs were collected for the cA_4_-bound complex and 6209 for the cA_4_-bound complex, respectively. Particles were picked using Topaz ^36^, followed by multiple rounds of 2D classification using CryoSPARC ^37^, resulting in 4,512,573 and 1,346,130 total particles for the two complexes, respectively.

For the TaqCad1-cA_4_ complex, hetero-refinement in CryoSPARC and several rounds of 3D refinement/classification using Relion 4.0 ^38^ without symmetry further led to 2,731,299 high-quality particles. A mask was constructed around each Cad1 dimer that was then used to extract dimer particles through symmetry expansion while subtracting unmasked signals (Supplementary Figure 3), leading to 12,364,827 good dimer particles that were 3D reconstructed. In parallel, the ADA domain portion was reconstructed by applying C3 symmetry of the intact particles. A composite map was made by applying the dimer density onto ADA map 3 times according to the C3 symmetry.

The apo TaqCad1 structure was constructed similarly, although without the focus on the CARF dimers. Good hexamer particles were selected for further refinement. Hetero-refinement in CryoSPARC and several rounds of 3D refinement/classification using Relion 4.0 ^38^ with C3 symmetry, followed by several rounds of non-uniform refinement ^39^ in CryoSPARC to reach the final 3D structures.

Structural models were built with manual adjustment in COOT ^40^ and refined in PHENIX ^41^ to satisfactory stereochemistry and real space map correlation parameters.

### UV absorbance analysis

TaqCad1-mediated reactions were assembled by incubating 2 μM TaqCad1 or its variants with or without 20 μM cA_4_ or cA_6_ (BIOLOG Life Science Institute) in a buffer containing 50 mM HEPES pH 7.5, 50 mM NaCl, 1 mM TCEP and 10 mM ZnCl_2_, or 1 mM MgCl_2_ or 1 mM CaCl_2_. The reactions were carried out at 55 °C for 30 min before being heat deactivated at 90 °C for 4 minutes and spun down at 13,2000 RPM for 20 minutes. UV absorbance of the reaction products were recorded on a NanoDrop spectrophotometer.

### HPLC analysis

High-performance liquid chromatography (HPLC) was employed to analyze ring nuclease activity and possible deamination products. For ring nuclease activity assay, reactions were initiated by incubating 5 μM TaqCad1 enzyme or its variants with 50 μM synthetic cA_4_ or cA_6_ (BIOLOG Life Science Institute) in a buffer containing 20 mM Tris-HCl pH 7.5, 150 mM NaCl, and 1 μM Zn^2+^. The reactions were carried out at 37 °C for 10–60 min before being heated at 95 °C for 10 minutes and spun down at 13,2000 RPM for 20 minutes.

For possible deamination activity, reactions were initiated by incubating 2 μM wildtype or mutant TaqCad1 enzymes, 1 mM ATP and/or 20 μM synthetic cA_4_ or cA_6_ in a 100 μl reaction volume containing 50 mM HEPES 7.5, 50 mM NaCl, 1 mM TCEP (tris(2-carboxyethyl)phosphine), 1 mM MgCl_2_, or 1 mM CaCl_2_ or 10 μM ZnCl_2_ at 55°C for 30 minutes. The reaction mixtures were quenched by heating at 90°C. Calf intestinal alkaline phosphatase (CIP) was added to dephosphorylate the products and followed by heat deactivation. The reaction mixtures were then filtered using Amicon® Ultra Centrifugal Filter with 3 kDa cut-off.

The supernatant of both reactions was analyzed using an HPLC system (Shimadzu Prominence LC-20) fitted with a SunFire C18 column (4.6 mm × 150 mm, 3.5 μm particle size). A 5 μl sample volume was injected and separated by a linear gradient of eluent A (20 mM ammonium bicarbonate) and eluent B (100% acetonitrile) at a flow rate of 0.3 ml/min over a total 22-min duration. The gradient conditions were as follows: a gradient of 2–30% B from 0 to 12 min, from a transition to 95% B for washing from 12.1 to 16 min, a constant 95% B concentration from 16.1 to 17.0 min, and finally, return to 2% B for equilibration from 17.1 to 22.0 min.

### Mass spectrometry analysis

For mass spectrometry analysis, reaction mixtures were prepared identically to those used for HPLC. Following urea denaturing, the supernatant was analyzed on an Agilent 6230 TOF-MS with the Agilent Mass Hunter Workstation Software TOF 6500 series in positive ion mode. Spectrum was analyzed using Agilent Mass Hunter Qualitative Analysis Navigator v.B.08 and visualized using GraphPad Prism.

### Microscale thermophoresis

The binding affinities of TaqCad1 and its variants with cOA were evaluated by microscale thermophoresis (MST) binding assay, using the instrument of NanoTemper Monolith NT.115 (NanoTemper Technologies, Munchen, Germany). To track the movement of TaqCad1 during the experiments, the protein was labeled using the his-tag labeling kit RED-tris-NTA (MO-L008, Nanotemper Technologies). The labeling was performed in PBST buffer, which also served as the assay buffer for MST experiments. After labeling, 50 nM TaqCad1 protein was incubated on ice for 60 mins with varying concentrations of cOA and then loaded into Monolith NT.115 MST standard-treated capillaries (MO-K022, NanoTemper Technologies). The measurements were carried out at 25°C using a Monolith NT.115 instrument with MO. control software, setting the light-emitting diode (LED) or excitation power at 40-80%, and MST medium power. Data analysis was performed using GraphPad Prism.

### In vitro RNA cleavage assay

The RNA cleavage assays were performed in a cleavage buffer containing 20 mM MES pH 6.5 and 5mM MnCl_2_. The reactions were performed at 60°C for 20 min and contained 100 nM Csx1 and 1μM target RNA (Supplementary Table 2) and varying concentrations of TaqCad1. The reactions were quenched using 2x formamide dye (95% formamide, 0.025% SDS, 0.025% bromophenol blue, 0.025% xylene cyanol FF, 0.5 mM EDTA). The reaction products were heated at 70 °C/3 min and separated by 7 M Urea, 15% polyacrylamidegel electrophoresis (PAGE) gels in 1x Tris/Borate/EDTA (TBE) running buffer and were visualized by staining with SYBR Gold II (Invitrogen) stain.

### In vivo plasmid challenge assays

To assess the effect of *in vivo* expression and activation of TaqCad1, electrocompetent *E. coli* BL21-AI were co-transformed with two plasmids prior to target/non-target plasmid challenge. The first one, pTsuTypeIII, carried several genes from *T. succinifaciens* for the formation of type III CRISPR-Cas complexes: *csm1* to *csm5*, *cas6*, and a minimal CRISPR array carrying one spacer sequence. These were individually placed under the control of T7 promoters (Supplementary Figure 9). The second plasmid, called a pEffector, encoded for one of the effectors used in the assay (either TaqCad1 or TsuCard1), constitutively expressed by a lacUV5 promoter (Supplementary Figure 9). On the target and non-target plasmids, a non-coding RNA was placed under the control of a trc promoter and lacO (Supplementary Figure 9) ^33^. In the case of the target plasmid, this non-coding RNA is complementary to the spacer on pTsuTypeIII.

The transformations were carried out in triplicate, where 100 ng of either target or non-target plasmid were electroporated into each strain. After electroporation, the cells were recovered in SOC medium for 1 hr at 37°C. For calculating the relative transformation efficiencies, ten-fold serial dilutions of the recovered cells were plated onto LB agar containing 34 µg/mL chloramphenicol, 50 µg/mL carbenicillin, 50 µg/mL kanamycin, 0.2% arabinose, and 0.5 mM IPTG. The plates were incubated overnight at 37°C, and transformation efficiencies were quantified. Data was analyzed and visualized using R 4.3.0, with statistical significance calculated by a one-sided unpaired Welch’s t-test.

For the liquid culture growth assay, the procedure for electroporation and recovery were done as previously described. After recovery, the cells were wash three times with LB medium. The cells were then diluted to OD_600_ = 0.05, and 150 µL of each dilution was added to a 96-well plate and covered with 50 µL mineral oil. The plate was incubated in a BioTek Epoch 2 Microplate Spectrophotometer for 22 hr at 37°C, with double-orbital shaking. Growth, measured by OD_600_, was recorded every 10 min. Data was visualized using Python 3.10.

## Author contributions

C.W., Y.Z., D.A.Y. and H.L. designed the experiments, C.W. and D.A.Y. expressed and purified TaqCad1 proteins; Y.Z. performed cryoEM analysis, C.W. and D.A.Y. performed MST and deamination assays, D.A.Y. and A. C.W. perform HPLC analysis, D.A.Y. performed RNA cleavage assays. H.H. performed mass spectroscopy analysis, C.W., Y.Z., D.A.Y. and H.L. wrote the manuscript. All authors edited the manuscript and provided insightful comments.

## Acknowledgments

Authors acknowledge the use of instruments at the Biological Science Imaging Resource (BSIR) supported by Florida State University, The Laboratory for BioMolecular Structure (LBMS) and the Pacific Northwest Center for Cryo-EM (PNCC). Authors thank X. Fu of BSIR for assistance in cryoEM data collection; G. Seo of the Institute of Molecular Biophysics Protein Expression Facility for providing facilities and resources for protein expression; S. Miller of the FSU Sequencing facility for assistance with Sanger sequencing; We also thank the staff at LBMS, especially Guobin Hu, and at PNCC, especially Nancy Meyer, for assistance with data collection.

We acknowledge the use of the BSIR instrumentations, including the Titan Krios, funded by NIH grant S10 RR025080, the Vitrobot and Solaris Plasma Cleaner, supported by S10 RR024564, and the BioQuantum/K3, acquired through U24 GM116788 funding. We are also thankful to SECM4 (The Southeastern Center for Microscopy of Macro Molecular Machines) for screening and data collection through NIH grant R24 GM145964.

## Data availability

The atomic coordinates of the cryo-EM structures of TaqCad1-cA_4_ and Apo TaqCad1 have been deposited in the Protein Data Bank under the identifiers ### and ### in the Electron Microscopy Data Bank under the entries ### and ### respectively.

